# Vitamin D deficiency during pregnancy and its associated factors among third trimester Malaysian pregnant women

**DOI:** 10.1101/616805

**Authors:** Fui Chee Woon, Yit Siew Chin, Intan Hakimah Ismail, Marijka Batterham, Amir Hamzah Abdul Latiff, Wan Ying Gan, Geeta Appannah, Siti Huzaifah Mohammed Hussien, Muliana Edi, Meng Lee Tan, Yoke Mun Chan

## Abstract

**Background:** Despite perennial sunshine, vitamin D deficiency is prevalent among Malaysian especially pregnant women.

**Objective:** To determine the vitamin D status and its associated factors among third trimester pregnant women attending government health clinics in Selangor and Kuala Lumpur, Malaysia.

**Methods:** Information on socio-demographic characteristics, obstetrical history, vitamin D intake, supplement use, and sun exposure were obtained through face-to-face interviews. Serum 25-hydroxyvitamin D concentration was measured and classified as deficient (< 30 nmol/L), insufficient (30-50 nmol/L), and sufficient (≥ 50 nmol/L).

**Results:** Of the 535 pregnant women recruited, 42.6% were vitamin D deficient. They consumed an average of 8.7 ± 6.7 μg of vitamin D daily. A total of 80.4% of the vitamin D were obtained from the food sources, while 19.6% were from dietary supplements. Fish and fish products showed the highest contribution to vitamin D intake (35.8%). The multivariate generalized linear mixed models, with clinic as a random effect, indicates that higher intake of vitamin D is associated with lower risk of vitamin D deficiency among pregnant women (OR = 0.96; 95% CI = 0.93-0.99). Non-Malay pregnant women had lower odds of having vitamin D deficiency (OR = 0.13; 95% CI = 0.04-0.37) compared to Malays. No associations were found between age, educational level, monthly household income, work status, gravidity, parity, pre-pregnancy body mass index, total hours of sun exposure, total percentage of body surface area, and sun exposure index per day with vitamin D deficiency.

**Conclusions:** Vitamin D deficiency is prevalent among Malaysian pregnant women. Considering the possible adverse obstetric and fetal outcomes of vitamin D deficiency during pregnancy, antenatal screening of vitamin D levels and nutrition education should be emphasised by taking into consideration ethnic differences.

## Introduction

Vitamin D, an essential fat-soluble vitamin or steroid prohormone, plays an important role in the regulation of calcium and phosphorus homeostasis and bone mineralization [1]. There are three main sources of vitamin D which include sunlight exposure, dietary sources, and supplement intake. Sunlight exposure is the primary source of vitamin D and is mainly influenced by environmental and personal factors such as seasons, geographic latitude, skin type, the percentage of body surface exposed to sunlight, and clothing [2,3]. Once ingested or produced by the body through skin exposure to the ultraviolet B radiation from the sun, vitamin D3 (cholecalciferol) is transported to the liver and is hydroxylated to 25-hydroxyvitamin D (25(OH)D) [4]. 25(OH)D is the major circulating form of vitamin D in human body [5]. Serum 25(OH)D is widely recognized as the best biochemical indicator of vitamin D status as it well-reflects the cumulative exposure to sunlight and dietary vitamin D intake of an individual [6]. Identifying the level of circulating 25(OH)D is important for diagnosis and monitoring of vitamin D deficiency [6].

Vitamin D deficiency has been identified as a global health problem and has affected more than 1 billion people globally [7,8], especially among pregnant women. The prevalence of vitamin D deficiency and insufficiency during pregnancy ranges from 27.0% to 91.0% in the United States, 39.0% to 65.0% in Canada, 45.0% to 100.0% in Asia, 19.0% to 96.0% in Europe, and 25.0% to 87.0% in Australia and New Zealand [8]. Despite being a tropical country with perennial sunshine, vitamin D deficiency in pregnant women has been reported in Malaysia. A recent study conducted at a tertiary hospital in Kuala Lumpur found that 71.7% of the third trimester pregnant women had vitamin D deficiency and 21.0% had vitamin D insufficiency [9]. Another local study reported 90.4% of the first trimester pregnant women in the Klang Valley had vitamin D insufficiency and deficiency [10]. A cohort study in Kelantan, Malaysia found that 59.8% and 37.3% of pregnant women had vitamin D deficiency during their second and third trimesters, respectively [11].

Low maternal vitamin D levels during pregnancy have been linked with multiple adverse obstetric outcomes such as maternal osteomalacia [12], gestational diabetes [13], preeclampsia [14], and primary cesarean section [15]. In addition, gestational vitamin D deficiency is associated with fetal intrauterine growth restriction and various adverse fetal and neonatal health outcomes, including higher risk of premature birth [16], abortion [17], low birth weight [18], neonatal hypocalcaemia [19], and childhood obesity [20].

Given the high prevalence of vitamin D deficiency among pregnant women and its adverse pregnancy outcomes, there is an urgent need to determine factors contributing to vitamin D deficiency during pregnancy in order to design effective prevention strategies that might reverse these alarming trends. Therefore, the aim of this study was to determine the prevalence of vitamin D deficiency among pregnant women in Selangor and Kuala Lumpur and to identify potential factors associated with vitamin D deficiency during the third trimester of pregnancy.

## Materials and Methods

### Ethics Statement

Ethical approvals for the study were obtained from the Ethics Committee for Research Involving Human Subjects, Universiti Putra Malaysia [FPSK(FR16)P006] and the Medical Research and Ethics Committee, Ministry of Health Malaysia (NMRR-16-1047-30685).

### Study design and respondents

This study is part of the Mother and Infant Cohort Study (MICOS) and the protocol of the study was previously described [21]. This study was conducted at six selected government Maternal and Child Health clinics in the state of Selangor and the city of Kuala Lumpur, Malaysia. Written informed consent was obtained from the respondents prior to data collection. Between November 2016 and January 2018, Malaysian women aged 18 years and above with singleton pregnancies of more than 28 weeks of gestations were invited to participate in the study during their routine prenatal check-ups at the selected clinics. Women with multiple pregnancies and planned to move out of the study area in the next one year were excluded from the study. Out of 3982 pregnant women who were invited to participate, 535 women consented and completed the study.

### Maternal vitamin D status

Vitamin D status was determined based on serum 25(OH)D analysis. A venous blood sample (2ml) was collected from the respondents and their serum 25(OH)D was measured by using the Siemens ADVIA Centaur^®^ Vitamin D Total assay (Siemens, Tarrytown, NY, USA). Serum 25(OH)D level was classified into vitamin D deficiency (< 30 nmol/L), vitamin D insufficiency (30-50 nmol/L) and vitamin D sufficient (≥ 50 nmol/L) [22].

### Maternal characteristics

Socio-demographic data including age, ethnicity, educational level, working status, monthly household income, and obstetrical history such as parity and gravidity were obtained from the respondents through a face-to-face interview. Pre-pregnancy body weight and height were obtained from medical records. Pre-pregnancy Body Mass Index (BMI) was calculated and classified based on World Health Organization (WHO) cut-off points [23].

### Maternal vitamin D intake and supplementation

Vitamin D intake and supplementation were assessed using a Vitamin D Food Frequency Questionnaire over the past month [24]. As vitamin D content is not available in Malaysian food composition table, the vitamin D content of raw food was obtained from the United States Department of Agriculture National Nutrient Database for Standard Reference [25] and Food Composition System Singapore, while vitamin D content of the commercial products were obtained from product labels. The daily average vitamin D intake (μg/day) was calculated by multiplying the frequency of consumption per day, serving size consumed, and vitamin D content of the food. The vitamin D intake was then compared with the Recommended Nutrient Intakes (RNI) for Malaysians [26] to determine the nutrient intake adequacy. The percentage contribution of each food group to total vitamin D intake was calculated to determine the main food sources of vitamin D.

### Maternal sun exposure

Sun exposure was assessed by using a Seven-day Sun Exposure Recall [27]. Respondents were required to record their outdoor activities over the past one week in terms of type of activity, duration (in minutes), frequency (per week), clothing, sunscreen use, gloves, and umbrellas. Body surface area (BSA) exposed to sunlight was estimated by using the “Rule of Nine” [27]. Sun exposure index (SEI) was calculated by multiplying the amount of time spent outdoors with BSA exposed.

### Data analysis

The IBM SPSS Statistics 24 software (SPSS Inc., Chicago, IL, USA) was used to analyse the data. Descriptive statistics such as mean and standard deviation (SD), as well as frequency and percentage were performed. Generalized linear mixed models (GLMM) were used to examine the associations between socio-demographic factors (gestation age, ethnicity, educational level, working status, monthly household income), obstetrical factors (gravidity, parity, pre-pregnancy BMI), and behavioral factors (vitamin D intake, intake of supplements contain vitamin D, total hours of sun exposure per day, total percentage of BSA per day, total SEI per day) with vitamin D deficiency during pregnancy. First, a model was fitted with only clinic entered as a random effect to determine the within-clinic intra-class correlation coefficient. Second, socio-demographic, obstetrical, and behavioral factors were individually added as fixed effects in the model adjusted for clinic clustering. Only variables that were significant at the 0.05 level were retained for the final model. Third, a final model was fitted with the socio-demographic, obstetrical, and behavioral factors that were found to be significantly associated with vitamin D deficiency, and associations among these variables were assessed while controlling for clinic clustering. Data were presented as odd ratios (OR) with 95% confidence interval (CI).

## Results

### Characteristics of the respondents

The mean serum 25(OH)D concentration for the total 535 pregnant women was 33.8 nmol/L (SD = 12.9) (Table 1). Based on the Institute of Medicine (IOM) classification [22], the prevalence of vitamin D deficiency, vitamin D insufficiency, and normal vitamin D was 42.6%, 49.3%, and 8.0%, respectively. Their mean age at conception was 29.9 (SD = 4.1) years. Majority of the respondents were Malay (92.1%), attained a tertiary education (81.7%), and had a moderate household income (52.3%). Most of them were employed (69.0%), multigravida (65.2%), and nulliparous (42.1%). In relation to pre-pregnancy BMI, the prevalence of underweight, overweight, and obesity was 9.0%, 25.0%, and 11.8%, respectively. The respondents consumed an average of 8.7 ± 6.7 μg of vitamin D daily, with three-quarters of them did not achieve the RNI for vitamin D which is 15 μg/day (74.4%). Overall, the median SEI of the respondents was 0.57. The median percentage of BSA of the respondents was 1.14% by taking into account of the face, neck, arms, hands, legs, and feet being exposed to the sunlight, as well as the clothing and the usage of sunscreen. The respondents spent about of 4.29 minutes per day being exposed to the sunlight.

**Table 1.** Characteristics of the respondents (n = 525).

| Characteristics |  | n | % |
| --- | --- | --- | --- |
| 25(OH)D (nmol/L) | Mean (SD) | 33.8 (12.9) |  |
|  | Deficient (< 30 nmol/L) | 228 | 42.6 |
|  | Insufficient (30-50 nmol/L) | 264 | 49.4 |
| | Sufficient (≥ 50 nmol/L) | 43 | 8.0 |
| Age at conception (years) | Mean (SD) | 29.9 (4.1) |  |
| Ethnicity | Malay | 439 | 92.1 |
|  | Non-Malay | 42 | 7.9 |
| Educational level | Secondary | 98 | 18.3 |
|  | Tertiary | 437 | 81.7 |
| Monthly household income<sup>a</sup> | Low (< RM2300) | 93 | 17.4 |
|  | Moderate (RM2300-RM5599) | 280 | 52.3 |
|  | High (> RM5600) | 162 | 30.3 |
| Work status | Non-working | 166 | 31.0 |
|  | Working | 369 | 69.0 |
| Gravidity | Primigravida | 186 | 34.8 |
|  | Multigravida | 349 | 65.2 |
| Parity | Nulliparous | 225 | 42.1 |
|  | Primiparous | 139 | 26.0 |
|  | Multiparous | 171 | 32.0 |

**Table 1.** (Continued)
| Characteristics |  | n | % |
| --- | --- | --- | --- |
| Pre-pregnancy BMI (kg/m<sup>2</sup>) | Mean (SD) | 24.1 (4.9) |  |
|  | Underweight (< 18.5 kg/m<sup>2</sup>) | 48 | 9.0 |
|  | Normal weight (18.5-24.9 kg/m<sup>2</sup>) | 290 | 54.0 |
|  | Overweight (25.0-29.9 kg/m<sup>2</sup>) | 134 | 25.0 |
|  | Obesity (≥ 30.0 kg/m<sup>2</sup>) | 63 | 11.8 |
| Dietary vitamin D intake (µg/day) | Mean (SD) | 10.2 (7.9) |  |
|  | Below RNI (< 15 µg/day) | 398 | 74.4 |
|  | Above RNI (≥15 µg/day) | 137 | 25.6 |
| Intake of supplements containing vitamin D | No | 355 | 66.4 |
|  | Yes | 180 | 33.6 |
| Total minutes of sun exposure per day | Median (IQR) | 4.29 (0.00, 17.14) |  |
| Total % BSA per day | Median (IQR) | 1.14 (0.00, 5.14) |  |
| SEI per day | Median (IQR) | 0.57 (0.00, 0.57) |  |

BSA, Body Surface Area; IQR, Interquartile Range; RM, Ringgit Malaysia; RNI, Recommended Nutrient Intakes;
SEI, Sun Exposure Index
<sup>a</sup>1 US dollar = RM 4.09 (as of March 16, 2019)

A total of 80.4% of the vitamin D were obtained from the food sources, while the rest were from dietary supplements (19.6%) (Table 2). Only one in three of the respondents took supplements containing vitamin D during pregnancy (33.6%). Fish and fish products (35.8%) showed the highest contribution to vitamin D intake, followed by milk and milk products (28.2%), eggs (9.1%), meat and meat products (3.9%), others (1.3%), beverages (1.2%), and cereal and cereal products (0.9%).

**Table 2.** Contribution of Food Items Towards the Daily Mean Intake of Vitamin D among the Respondents.

| Food item | Contribution (%) |
| --- | --- |
| Fish and fish products | 35.87 |
| Indian mackerel | 13.78 |
| Eastern little tuna | 8.90 |
| Prawn | 4.59 |
| Spanish mackerel | 4.51 |
| Salmon | 2.11 |
| Anchovy | 0.98 |
| Canned sardine | 0.93 |
| Canned tuna | 0.05 |
| Canned mackerel | 0.02 |
| Milk and milk products | 28.19 |
| Fresh milk (Full cream/Low-fat/Flavored) | 19.31 |
| Maternal milk powder | 4.82 |
| Milk powder (Full cream/Low-fat) | 2.71 |
| Sweetened condensed milk | 0.49 |
| Cheese | 0.41 |
| Ice-cream | 0.25 |
| Butter | 0.20 |
| Eggs | 9.13 |
| Meat and meat products | 3.85 |
| Chicken | 3.54 |

**Table 2.** (Continued)
| Food item | Contribution (%) |
| --- | --- |
| Beef | 0.20 |
| Beef sausage | 0.05 |
| Pork | 0.04 |
| Cow liver | 0.02 |
| Others | 1.31 |
| Margarine | 0.88 |
| Mushroom | 0.38 |
| Mashed potatoes | 0.05 |
| Beverages | 1.22 |
| Cultured milk drinks | 1.21 |
| Fortified soy drinks | 0.01 |
| Cereal and cereal products | 0.86 |
| Cereal drinks | 0.83 |
| Pancake | 0.02 |
| Waffle | 0.01 |
| Supplements containing vitamin D | 19.57 |

### Factors associated with maternal vitamin D deficiency

As shown in Table 3, the estimated intercept and 95% CI in the null model (Model 1) was 0.01 (95% CI = 0.00, 1.50). In the bivariate model adjusted for clinic clustering (Model 2), non-Malay (OR = 0.13, 95% CI = 0.04, 0.37), intake of supplements containing vitamin D (OR = 0.52, 95% CI = 0.36, 0.75), and higher dietary vitamin D intake (OR = 0.96, 95% CI = 0.93, 0.98) were significantly associated with lower risk of vitamin D deficiency compared to their counterparts. No associations were found between age, educational level, monthly household income, work status, gravidity, parity, pre-pregnancy BMI, total hours of sun exposure, total percentage of BSA, and SEI per day with vitamin D deficiency.

**Table 3.**
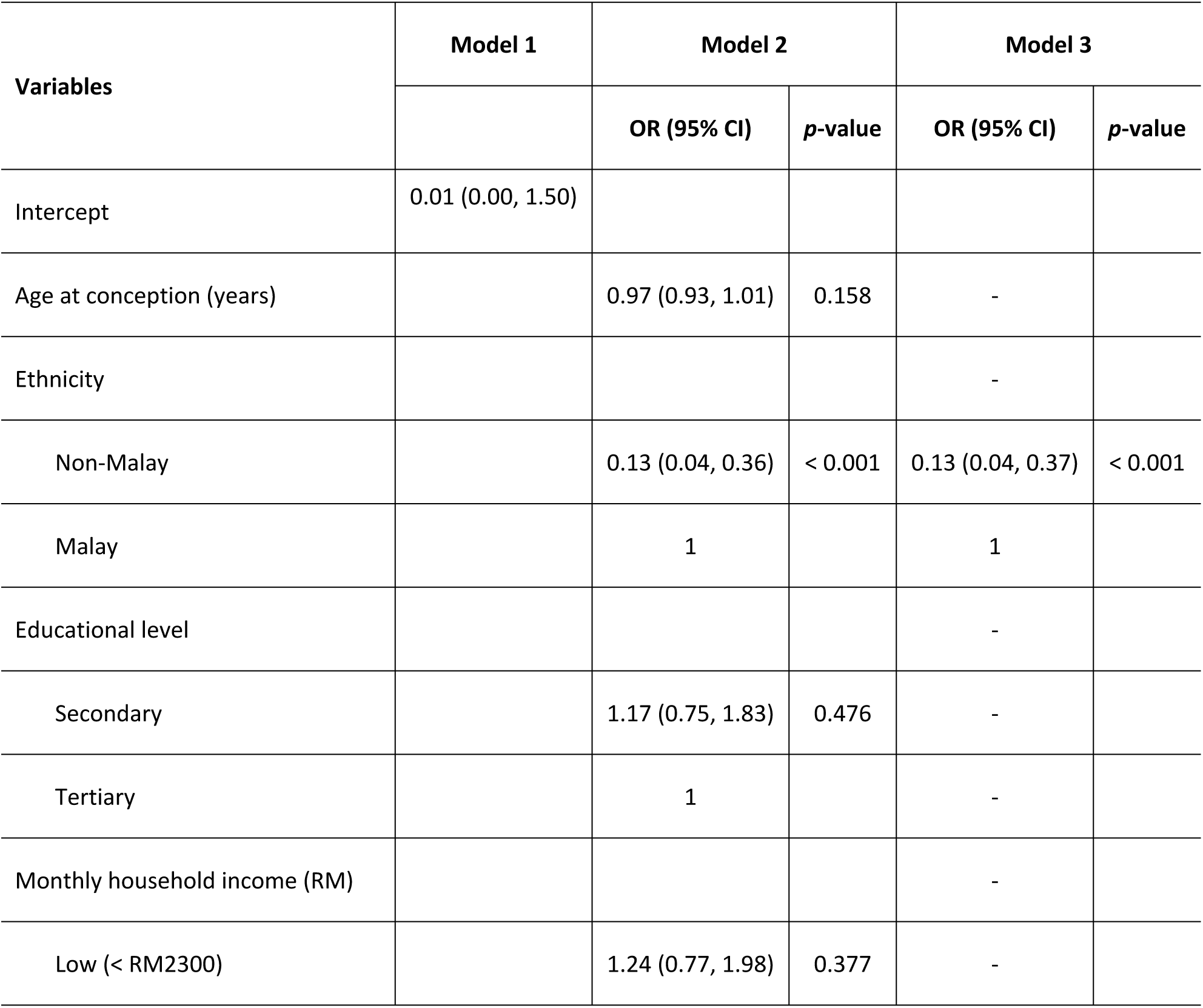
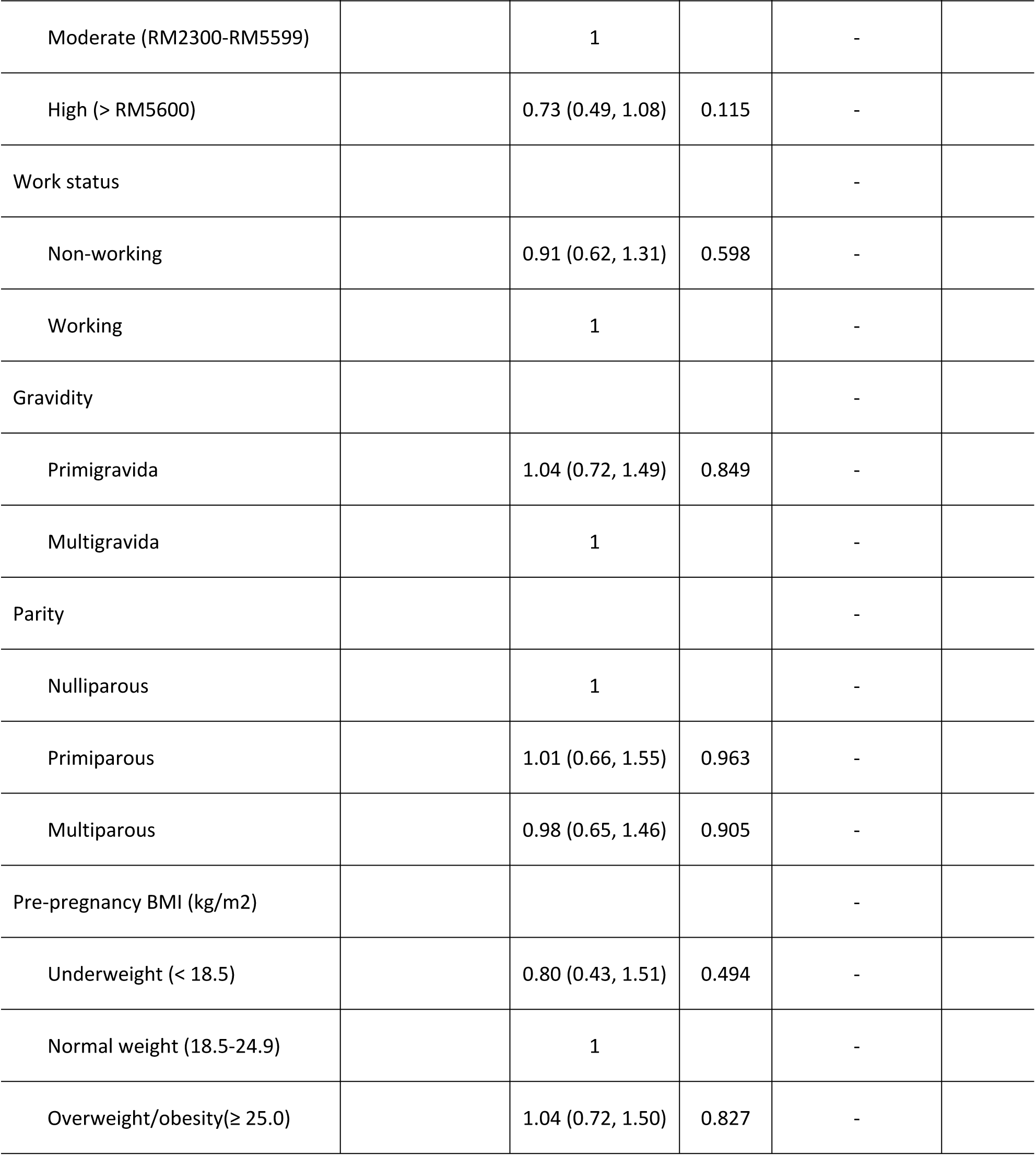
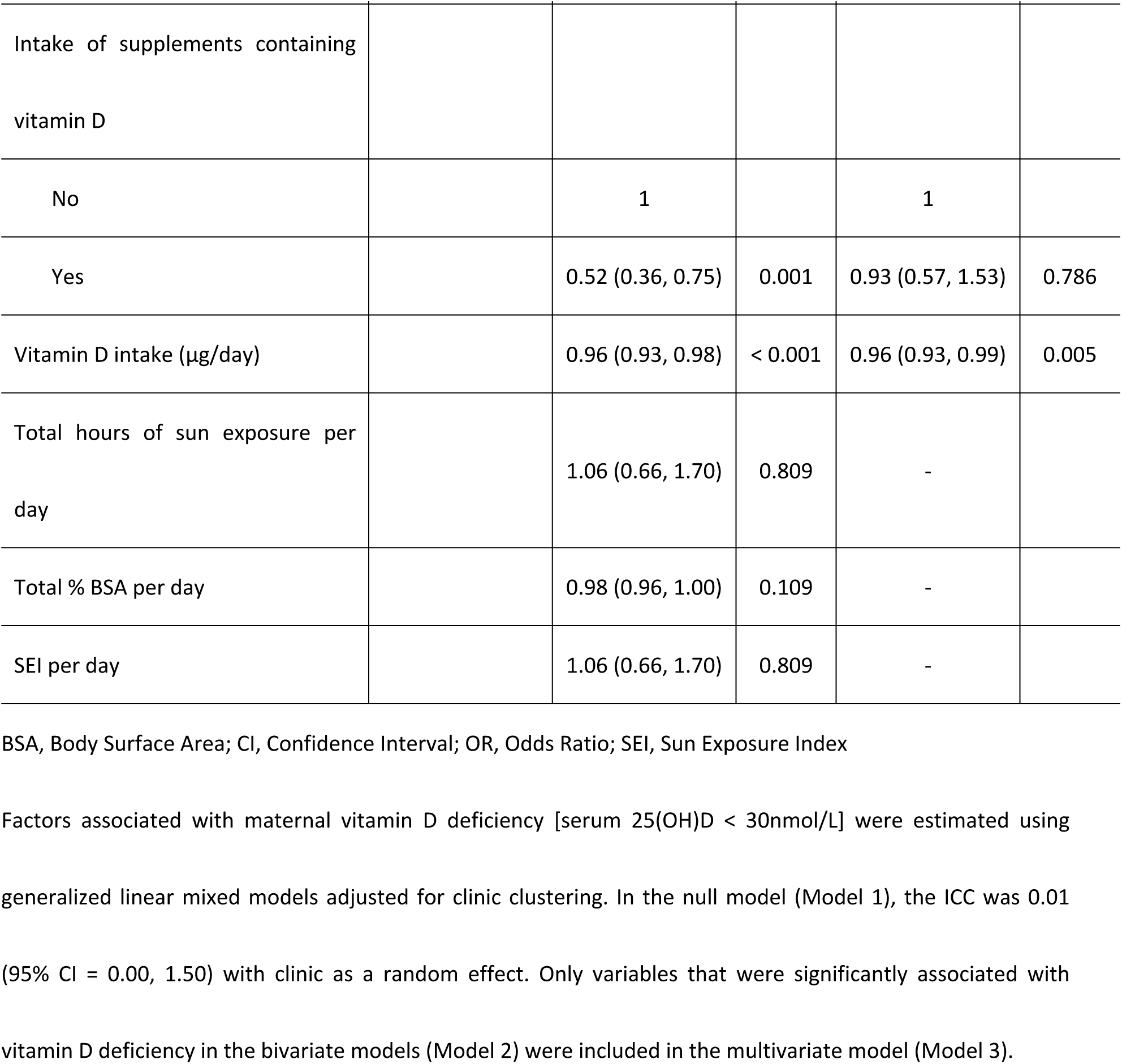
Factors Associated with Maternal Vitamin D Deficiency [25(OH)D <30 nmol/L].

In the multivariate model adjusted for clinic clustering (Model 3), ethnicity and dietary vitamin D intake remained significant. The odds of having vitamin D deficiency among non-Malays were 0.13 times lower than Malays (OR = 0.13, 95% CI = 0.04, 0.37). Meanwhile, pregnant women who had higher intake of vitamin D were less likely to have vitamin D deficiency during pregnancy (OR = 0.96, 95% CI = 0.93, 0.99). The association between intake of supplements containing vitamin D with vitamin D deficiency was no longer significant.

## Discussion

The present study revealed that 42.6% of the pregnant women were vitamin D deficiency and almost half were vitamin D insufficiency (49.3%). Women who had higher intake of dietary vitamin D and being non-Malays were less likely to have vitamin D deficiency during pregnancy.

High prevalence of vitamin D deficiency and insufficiency have been reported in several studies in the tropical countries [28-30]. A recent study conducted in West Sumatra, Indonesia reported the prevalence of vitamin D deficiency and insufficiency among third trimester pregnant women was 61.3% [28]. Another study found that 60.0% of Vietnamese women at 32 weeks gestation had low vitamin D levels [29]. In Thailand, 75.5% of the pregnant women had hypovitaminosis at the time of giving birth [30]. The prevalence of vitamin D deficiency and insufficiency in the present study was much higher than those reported in the aforementioned studies which used different serum 25(OH)D cut-off level of < 75 nmol/L. To date, there is still lack of consensus on the definition of vitamin D levels. While IOM defined a serum 25(OH)D level less than 30 nmol/L as deficiency and 30-50 nmol/L as insufficiency [22], the Endocrine Society Task Force set a higher cut-off values for vitamin D deficiency [25(OH)D < 50 nmol/L] and insufficiency [25(OH)D 50-74 nmol/L] [31]. The IOM definitions were used in this study as findings from previous studies indicated that a deficient serum 25(OH)D level below 30 nmol/L was associated with increased risk of adverse skeletal health outcomes including osteomalacia [22]. Meanwhile, insufficient serum 25(OH)D level of 30-50 nmol/L could lead to hyperparathyroidism, accelerated bone turnover and osteoporosis [22].

In this study, pregnant women who had higher intake vitamin D were more likely to have lower risk of vitamin D deficiency. This finding is in agreement with Shiraishi et al. [32] that found higher vitamin D intake significantly contributed to higher serum 25(OH)D concentration among pregnant women. This could be attributed to the high consumption of vitamin D containing food such as milk and milk products and fish and fish products (as shown in Table 3). Similarly, a recent local study conducted by Yong et al. [33] demonstrated that milk and dairy products were the major food sources contributing to vitamin D intake among pregnant women.

In the current study, we found that among third trimester pregnant women, those who were Malays were at a higher risk for vitamin D deficiency as compared to the non-Malays. The significant ethnic differences in the prevalence of vitamin D deficiency was in line with previous studies conducted among general population and pregnant women in Malaysia, showing that Malays had the highest prevalence of vitamin D deficiency than non-Malays [19,34]. The high prevalence of vitamin D deficiency might be due to religious and cultural reasons. Muslim women are compulsory to cover entire body parts [35] and this reduces the probability for the Malay pregnant women to get sufficient sunlight, which will then lower the vitamin D production in their body. Similarly, previous studies conducted in Islamic countries such as Iran and Pakistan reported high prevalence of vitamin D insufficiency and deficiency among Muslim pregnant women [36,37].

Only one in three women in the study were taking dietary supplements containing vitamin D, such as multivitamins and calcium supplements enriched with vitamin D. Intake of supplements containing vitamin D significantly lowered the risk of vitamin D deficiency in the bivariate model but was no longer significant in the multivariate model. This finding was inconsistent with previous studies conducted among pregnant Japanese [32] and Chinese [38] women, in which the use of vitamin D supplements and multivitamins were associated with higher serum 25(OH)D levels. One of the possible explanations for these findings is that the use of vitamin D supplements was uncommon among Malaysian pregnant women. We also found that the major contributor of vitamin D was from food sources, while dietary supplements only contributed towards less than a quarter of the total vitamin D intake.

In line with the findings reported in a local study conducted among pregnant women in an urban district in Malaysia [10], no association was found between sun exposure and vitamin D levels in this study. This might be due to low sun exposure in this population whereby majority of them (73.5%) spent less than 10 minutes in a week between 10am to 2pm in the afternoon when vitamin D from the sunlight is synthesized most efficiently by the body (unpublished results).

This study has several limitations. First, the cause-effect relationships between factors and vitamin D deficiency cannot be determined from the cross-sectional study design. Second, self-reported data on sun exposure and dietary vitamin D intake may lead to recall bias. High proportion of the respondents were Malays while only 7.9% were from other ethnic groups such as Chinese and Indian. Thus, the present results could not be generalized to all pregnant women in Malaysia. We acknowledge that other potential factors which may contribute to vitamin D levels, such as skin type, physical activity, season, or genetic background, were not examined in the present study and warrant further studies.

## Conclusions

Although Malaysia is a country with abundant sunshine all year round, vitamin D deficiency was highly prevalent among third trimester pregnant women. High intake of vitamin D was found to be a protective factor for vitamin D deficiency, while Malay women had a higher risk of vitamin D deficiency. Future interventions for the prevention and control of maternal vitamin D deficiency should take into account of the ethnic differences. Considering the long term health complications of vitamin D deficiency during pregnancy, antenatal screening of vitamin D levels and nutrition education should be emphasized among pregnant women.

## Acknowledgments

The authors would like to acknowledge the research enumerators, medical assistants, and MCH nurses for their assistance and contribution in this study.

## Author Contributions

Conceptualization: FCW YSC IHI YMC AHAL WYG GA. Methodology: FCW YSC IHI YMC MB AHAL WYG GA. Formal analysis: FCW YSC MB SHMH ME MLT. Investigation: FCW SHMH ME MLT. Data curation: FCW YSC SHMH ME MLT. Writing - original draft preparation: FCW. Writing - review and editing: FCW YSC IHI YMC MB AHAL WYG GA SHMH ME MLT. Supervision: YSC. Funding acquisition: FCW YSC.

## Supporting information

**S1 Dataset.** Dataset for study on vitamin D deficiency during pregnancy and its associated factors among third trimester Malaysian pregnant women (n=535).

